# A biomimetic fruit fly robot for studying the neuromechanics of legged locomotion

**DOI:** 10.1101/2024.02.22.581436

**Authors:** Clarus A. Goldsmith, Moritz Haustein, Ansgar Büschges, Nicholas S. Szczecinski

**Author notes:** This paper has supplementary downloadable material available at http://ieeexplore.ieee.org, provided by the authors. This includes two multimedia MP4 format movie clips, which show the robot walking.

## Abstract

For decades, the field of biologically inspired robotics has leveraged insights from animal locomotion to improve the walking ability of legged robots. Recently, “biomimetic” robots have been developed to model how specific animals walk. By prioritizing biological accuracy to the target organism rather than the application of general principles from biology, these robots can be used to develop detailed biological hypotheses for animal experiments, ultimately improving our understanding of the biological control of legs while improving technical solutions. In this work, we report the development and validation of the robot Drosophibot II, a meso-scale robotic model of an adult fruit fly, *Drosophila melanogaster*. This robot is novel for its close attention to the kinematics and dynamics of *Drosophila*, an increasingly important model of legged locomotion. Each leg’s proportions and degrees of freedom have been modeled after *Drosophila* 3D pose estimation data. We developed a program to automatically solve the inverse kinematics necessary for walking and solve the inverse dynamics necessary for mechatronic design. By applying this solver to a fly-scale body structure, we demonstrate that the robot’s dynamics fits those modeled for the fly. We validate the robot’s ability to walk forward and backward via open-loop straight line walking with biologically inspired foot trajectories. This robot will be used to test biologically inspired walking controllers informed by the morphology and dynamics of the insect nervous system, which will increase our understanding of how the nervous system controls legged locomotion.

## I. Introduction

**W**HILE scientists and engineers have increasingly realized the potential of robots to complete tasks in real-world environments, the design and control of legs has been a persistent topic of interest. Robots with legs can traverse natural and man-made terrains that are non-traversable with wheels or treads. However, their additional degrees of freedom (DoF) make them more mechanically complex and difficult to control. For the past several decades, the field of biologically inspired robotics has used animal locomotion as a template to improve the capability of robots [for reviews, see [1], [2], [3], [4]]. The structures of animal legs have evolved over hundreds of millions of years to traverse dynamic and uncertain terrains. Consequently, animal nervous systems are finely tuned to control these structures in a robust and adaptable manner. By applying principles from animal neuromechanics to robots, engineers are able to endow robots with similar capabilities and improve their locomotion. Such an approach has been successfully applied to many modes of locomotion including swimming [5], [6], [7], climbing [8], [9], [10], and flying [11], [12], [13] in addition to walking [14], [15], [16], [17].

More recently, a new, “biology-first” approach to designing biomimetic robots has developed to model how specific animals walk [18], [19], [20], [21], [22], [23], [24]. By closely modeling the neuromechanics of their target animal, these “biomimetic” robots serve as testbeds for neuromechanical hypotheses. Such an approach advances our understanding of how the nervous system controls walking in addition to enhancing robotic walking capability.

Insects have consistently been a target for bio-inspired and biomimetic legged robotics [reviews in [25], [26], [27], [28]]. Despite the small size of their bodies and nervous systems, insects demonstrate a range of legged behaviors rivaling that of vertebrates. A variety of tools have been developed over the last 50+ years to investigate invertebrate nervous systems, providing a library of neurobiological insect data from which to develop robot controllers [29], [30], [31]. The hexapod structure of insects also allows for statically stable locomotion at all possible speeds without redundant legs, making them appealing models for robots [32], [33], [34]. Early insect-inspired robots near the end of the twentieth century leveraged these aspects of insects to produce robust walking through distributed controller architectures [14], [35], [36]. These early endeavors highlighted the benefits of a biologically-inspired approach in insect robots. As computational power advanced, similar insect robots began modeling the dynamics of neurons and synapses in their controllers to produce further capability [37], [38], [39].

As a result of the influx of biological data, a growing number of highly biomimetic insect robots have begun to be developed. These robots are designed with close consideration to both the nervous system and mechanics of their target animal. One example is the hexapod robot HECTOR, which was modeled after the stick insect, *Carausius morosus*. HECTOR’s design uses the relative distances of the leg attachment points, the orientation of leg joint axes, and the division into three body segments to inform the mechatronic design [22], [40]. The robot has been used to explore several mechanical aspects of bioinspired locomotion such as compliant actuation and distributed proprioception, as well as serving as the testbed for the decentralized walking controller, Walknet [39]. Another example is ALPHA, a robot modeled after the African ball-rolling dung beetle *Scarabaeinae galenus*. ALPHA utilized Micro-CT scans of dead specimens to precisely proportion the legs and body [23]. Joint axes’ positions and angles were also derived from these scans based on a series of cylinders and cones aligned with each joint’s surface curvature. The researchers behind ALPHA demonstrated several mechanical benefits of the robot’s biomimetic morphology compared to a traditional hexapod configuration, such as increasing possible step lengths and decreasing motor accelerations. These results provide motivation for copying the complexities of insect leg kinematics.

Although many insect species have been studied to unravel motor control of locomotion until today, the fruit fly, *Drosophila melanogaster*, has gained more and more popularity in the field of neurobiology over the last decades due to its tremendous potential to link behavior with the anatomy and physiology of the nervous system. There is accumulating knowledge about the anatomy of the brain and ventral nerve cord including the connectome [41], [42], [43]. The ever-growing genetic toolboxes available [for review: [44], [45]] for *Drosophila* also enable recording neuronal activity in behaving animals [46] and to manipulate the activity of specific neurons in a straight-forward manner [47], [48], [49], [50]. Further, several *Drosophila* studies have identified neurons similar to those found in other insects [51], [52], demonstrating the potential for data from other insects to be applied to *Drosophila*, and vice versa. All together these developments render *Drosophila* as a promising target organism for biomimetic insect robot construction.

Drosophibot [24], a meso-scale robot modeled after *Drosophila melanogaster*, was previously developed to further facilitate the testing of synthetic nervous system (SNS) controllers based on the insect nervous system [20], [53], [54], [55]. Drosophibot included several features to capture the animal’s biomechanics such as biomimetic actuator control and sensing and parallel elastic joints. The robot was developed before 3D motion capture data for the fly was available, so leg proportions and joint axes were instead approximated from video footage and inspired by other insects such as *C. morosus*. Each leg was manufactured identically to simplify mechatronic design. As a result, several of the joint axes and leg attachment points were not accurate to the animal, so the robot struggled to execute fly-like walking.

Now that more neuromechanical data is available for *Drosophila* and recent advances in pose estimation algorithms allow the recording of 3D kinematic data at the leg joint level [56], [57], the design of Drosophibot can be updated to more closely resemble the insect. In particular, we want the robot and the animal to be kinematically and dynamically similar. Kinematic similarity allows the robot’s legs to execute similar motions as the insect, which is crucial for investigating aspects of insect walking such as interleg coordination [58], [59], [34].

Dynamic similarity prevents a brain-body mismatch while using SNS controllers for neuromechanical investigations. *Drosophila*’s nervous system has evolved over millennia to control walking in a specific leg structure with a corresponding combination of inertial, gravitational, elastic, and viscous forces acting within the legs. The interplay of these types of forces, which are tied to the animal’s size and walking speed, has been found to significantly influence how the nervous system controls walking [60], [61], [62]. A SNS developed from *Drosophila* data will be naturally tuned to produce this type of control, so making the robot dynamically similar despite its larger size prevents incongruities with the controller. Dynamic similarity will also allow us to investigate biomimetic sensory feedback during walking from sense organs such as the campaniform sensilla (CS), mechanoreceptors that encode exoskeleton strain [63], [64].

### A. Contributions

We present Drosophibot II, a biomimetic meso-scale robot modeled after the adult fruit fly, *Drosophila melanogaster*, to model and integrate the recent influx of *Drosophila* neurome-chanical data (Fig. 1). Drosophibot II’s leg proportions and degrees of freedom were closely modeled after 3D motion capture data from the insect performing straight line walking on a spherical treadmill. To our knowledge, it is the first biomimetic insect robot to utilize such data to inform its design. To design and validate the robot, we developed a program to generate biomimetic footpaths given key stepping parameters, solve the inverse kinematics necessary for walking, and calculate the inverse dynamics necessary to inform mechatronic design. We use this program to compare the joint angles and torques between the robot and a scale model of *Drosophila* and show that they are dynamically similar with and without parallel elastic components. We believe we are the first to apply such a rigorous consideration of dynamic similarity to the development of a biomimetic robot. The robot can successfully perform straight-line forward and backward walking. We also tested the robot at three different walking speeds to demonstrate that its walking is quasi-static, meaning that inertial forces do not impact its dynamics. This means that the non-biological mass distribution caused by the weight of the actuators in the legs does not meaningfully change the dynamics across walking speeds. In the future, this robot can be used to test walking controllers based on the neural control architecture of the insect nervous system, increasing our understanding of how the nervous system controls legs.

**Fig. 1.**
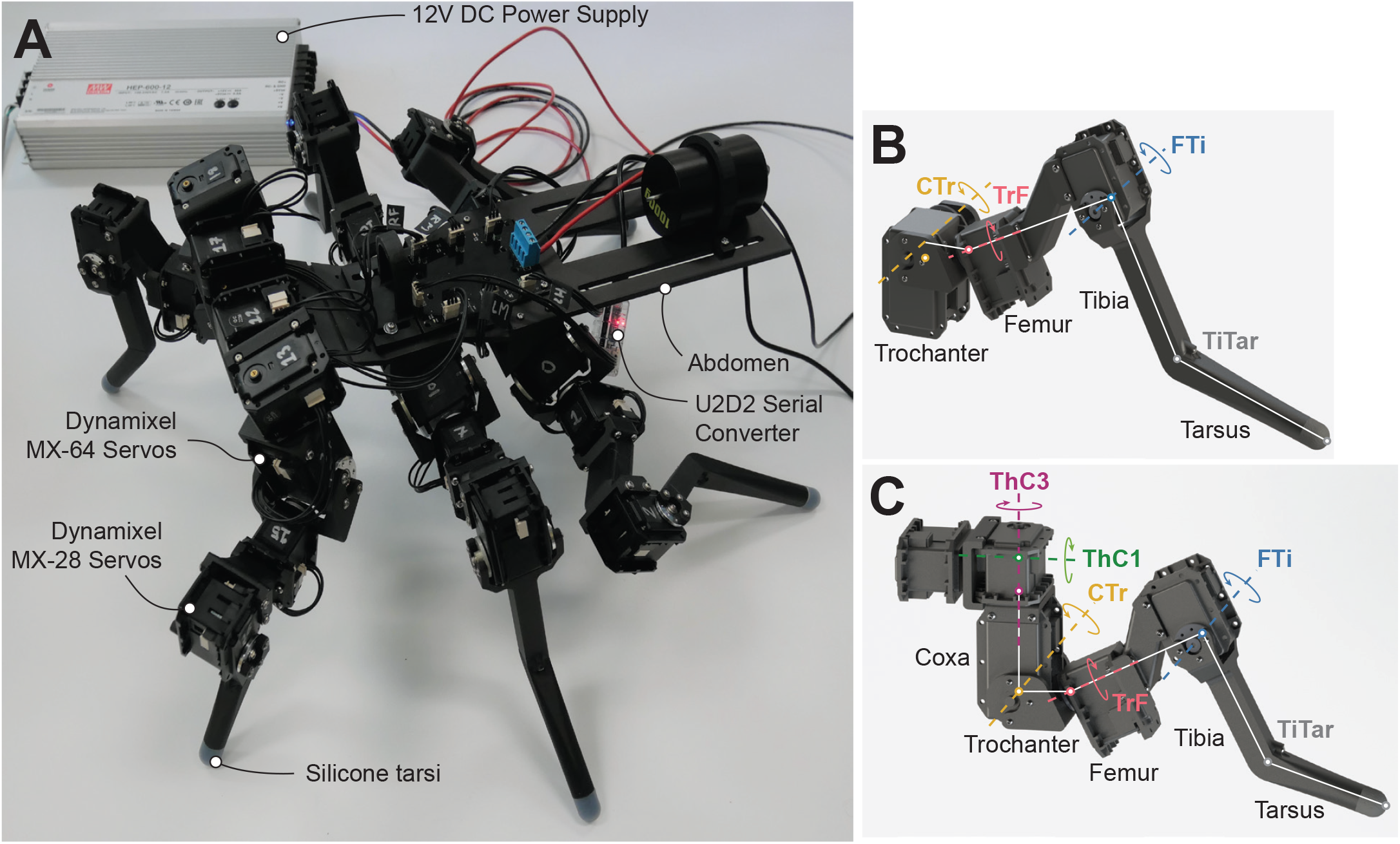
Drosophibot II, a biomimetic robot modeled after Drosophila melanogaster. (A) Photo of the robot with notable components annotated. (B) CAD drawing of a middle leg with the leg segments, joints, and axes of rotation labeled. The middle and hind legs have the same DoF and leg segments, though some segment lengths differ. (C) CAD drawing of a front leg with the leg segments, joints and axes of rotation labeled.

## II. Materials and Methods

### A. Fruit fly kinematics analysis

In order to closely characterize the kinematics of *Drosophila* on a robotic platform, we recorded tethered flies walking on a spherical treadmill (Fig. 2), then used DeepLabCut (DLC) [57] to automatically annotate the recordings and produce 3D pose estimation data for each trial. Appendix A describes this experimental setup and data collection in detail.

**Fig. 2.**
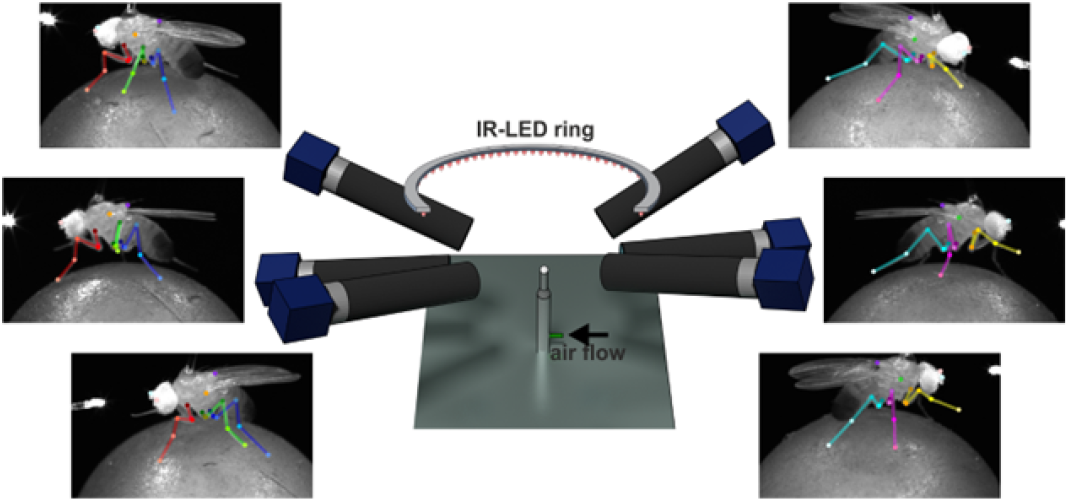
3D motion capture setup. Tethered flies walked on a spherical treadmill and leg movements were recorded with six high-speed cameras. Positions of leg and body features were tracked with DLC for subsequent 3D reconstruction.

The pose estimation data tracks the spatial location of each joint in the legs (a.k.a “joint-points”), as well as additional body parts required to transform the positional data into a body-centered coordinate system. We then used this data to analyze the kinematics of the different leg pairs. In particular, we were interested in identifying the minimum number of DoF that best approximate the movement in each of the fly’s leg pairs, as well as finding the average proportions between the different leg segments and leg attachment point distances on the body (e.g., the lateral distance between each pair of thorax-coxa (ThC) joints). By minimizing the number of actuated joints on the robot, we are able to minimize its overall actuator weight and increase its strength-to-weight ratio while enabling it to produce motions similar to those of the animal (e.g., walking in a line). Obtaining average leg and body ratios enabled us to construct the robot with “typical” fly proportions (i.e., kinematic similarity).

Our method for the analysis of leg DoF is presented in detail in [65] where it was applied to the middle and hind leg pairs. To briefly summarize the process, we begin by constructing a mathematical kinematic leg chain for every frame of motion capture data, normalized such that the ThC is the origin of the leg’s spatial frame. The leg chain contains seven total DoF that have been directly recorded or hypothesized in the animal: ThC protraction/retraction (ThC1), levation/depression (ThC2), and rotation (ThC3); coxa-trochanter (CTr) flexion/extension; trochanter-femur (TrF) rotation (TrF1) and flexion/extension (TrF2) ; and femur-tibia (FTi) flexion/extension [66], [67], [68], [69]. We use product of exponentials [70] to generate the leg chain with segment lengths calculated anew for each frame. We then find the “best fit” configuration of the model by solving for the joint angles that minimize the sum of Euclidean distances between each joint-point in the model and on the animal. After completing this process for the “complete” leg chain, various combinations of DoF were fixed at their average angle across an experimental trial to observe the effects these DoF have on the fit to the animal data. In that regard, we analyzed the average error in Euclidean distance of each joint-point as well as the average orientation and overall range of motion (RoM) of the “leg plane” (i.e., the plane containing the femur and tibia major axes). Orientation of the leg plane dictates the degree that moments from the ground reaction forces (GRF) are counteracted by the actuator versus passively resisted by the joint structure. Thus, considering the leg plane angle allowed us to consider biological force distributions.

Based on this method of analysis, we selected the CTr, TrF1, and FTi DoF as the mobile joints in our robot’s middle and hind leg pairs [65]. We then conducted analysis of the front limbs, as presented in Figure 3. In addition to minimizing the number of actuators, for the front limbs we were interested in DoF combinations that would: 1) Minimize the DoF in the ThC and 2) Include the TrF1. The former criterion would help minimize the mechatronic complexity of the ThC on the robot, as fewer actuators would need to fit into the limited space. The latter criterion allows us to investigate the role of the TrF1 DoF in leg pronation/supination, as we have in the middle and hind limbs. In regards to the mean error of each joint-point (Fig. 3A), fixing only the ThC2 produced the least error across all joints. The mean error of the leg plane angle to the vertical (Fig. 3B) for this combination of fixed DoF was on average 8 degrees higher than in the animal. However, a similar level of error occurred in every DoF combination tested. The RoM of the leg plane was higher than that in the animal, but was considered inconsequential. Based on this analysis, we included the ThC1, ThC3, CTr, TrF, and FTi DoF in Drosophibot II’s front legs, for five total DoF.

**Fig. 3.**
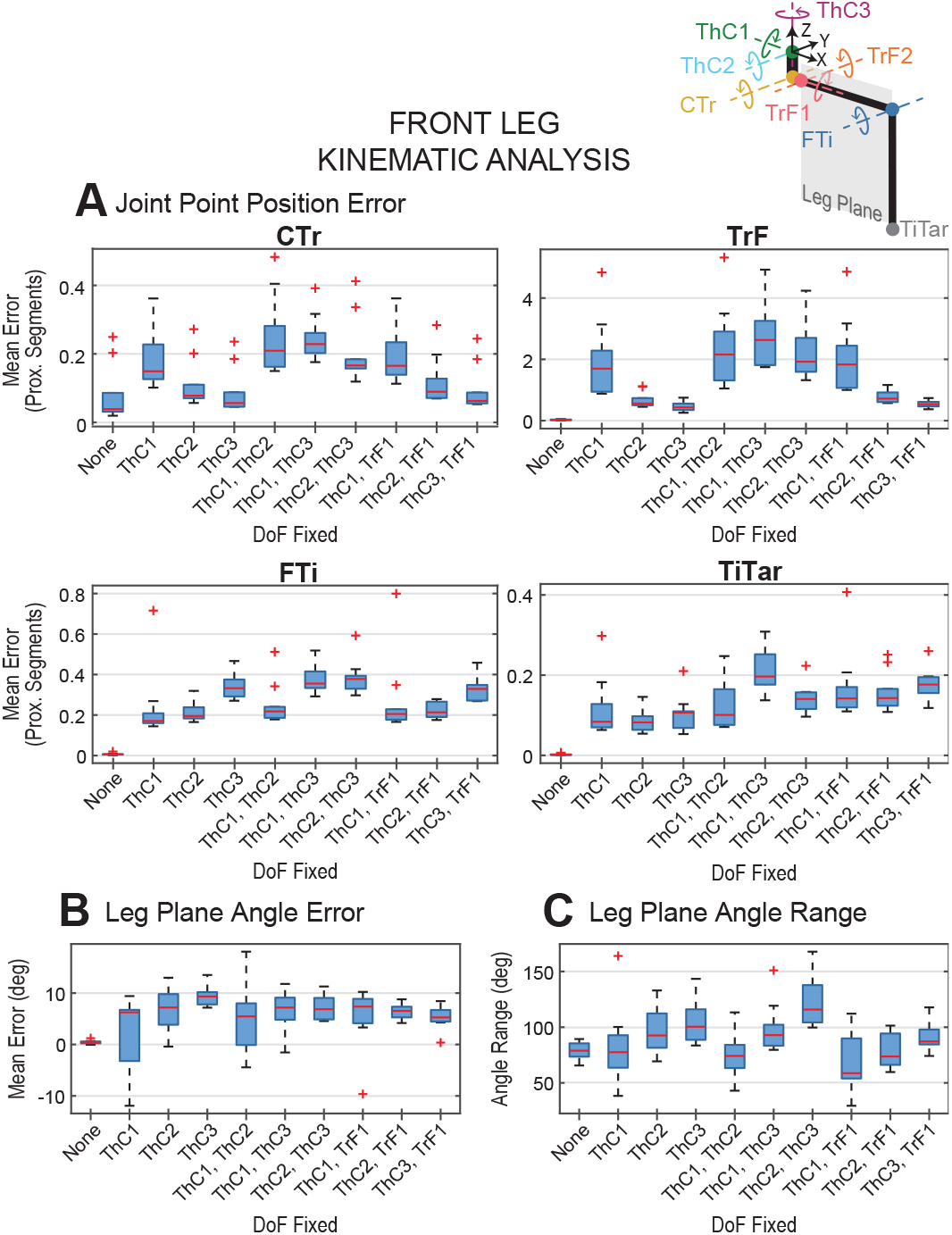
Kinematic analysis of the front limbs of Drosophila. Analysis of different fit parameters for a simulated leg chain attempting to match movements from Drosophila 3D motion capture data, as described in [65], with different DoF fixed at their average positions over a fully mobile trial. For all DoF fixation combinations except ‘None,’ the TrF2 DoF is also fixed. (A) Mean error of the spatial position of tracked points, corresponding to joint locations, over a trial. The error is normalized in terms of proximal segments (e.g., TrF error is in terms of trochanter length). (B) Mean error of the leg plane (i.e., the plane containing the femur and tibia major axes) over a trial. (C) Range of leg plane angles over a trial.

In addition to the DoF kinematic analysis, we determined the average proportions of the leg segments and body dimensions from our recorded flies (N=4/3 females/males). We first calculated the distances between the joint-points for each frame of video capture to estimate leg segment lengths. These values were averaged over a trial to determine approximate lengths in each fly, then averaged again to calculate the segment lengths of a “prototypical” fly. We scaled our robot such that the length of a middle leg femur was 10 cm. As such, we used our “prototypical” femur length in the fly data to determine length ratios between the femur and the other segments. Using these ratios, we calculated the leg segment lengths necessary for the robot to be proportional to the fruit fly. With our desired scaling constraint, Drosophibot II is approx. 140:1 scale to the insect.

### B. Inverse kinematic and dynamic solver

Using the leg and body proportions and the selected DoF from the kinematic analysis, we then created an inverse kinematic and dynamic solver in MATLAB (MathWorks, Natick, MA). The primary purposes of this solver were to generate robot limb trajectories given the body’s motion through space and calculate the joint torques, internal forces, and GRF in the robot during walking. Limb trajectories (e.g., joint angles necessary to move the foot in a particular way) can be used to control the robot, while aspects of the dynamics would inform actuator selection, power supply selection, and component design.

The solver begins by extracting position, dimension, and mass data (i.e., joint locations, axes, and limits; segment lengths, center of mass (CoM) locations, and masses) for each component of an organism’s structure from a user provided Animatlab [71] file. We use “organism” as a more general term to describe a given structure, as the solver may also be used to model animals. Using this provided structure, the solver generates a resting posture for the organism. Specifically, it attempts to set each joint’s angle close to the middle of its limits such that the foot’s vertical (y) component is at a user defined ground level and that the lateral (z) and sagittal (x) components satisfy positional constraints in relation to the leg’s ThC location adapted from the *Drosophila* stepping data [34]. Once a resting posture is found, the solver utilizes user-defined stepping parameters of walking speed, body translation and rotation per step, swing duration, and duty cycle to design stance-phase footpaths in each leg. The resting posture is used as the mid-point of these footpaths. A full list of the required parameters, as well as the values used to generate the data in this manuscript, can be found in Table I. Several of these parameters have previously been shown to be sufficient to predict biomimetic footpaths based on the desired movement of the body [34]. Using these footpaths, swing phase foot trajectories are calculated for the x, y, and z components separately as 5th order polynomials. For the x and z components, one 5th order polynomial is generated between the start and end of stance. For the y component, two polynomials are used; one interpolated between the start of stance and a user specified peak swing height, and the other between the swing peak and the end of stance. Currently, the horizontal location of the peak swing height is set to the middle of the stance footpath.

**TABLE I.** Walking parameters of Drosophibot II. These values result in total step period of between 1s-4s. Body height is varied between leg pairs while physically running the robot to counteract servo backlash and component deformation.

| Param. | Description | Value |  |  |
| --- | --- | --- | --- | --- |
|  |  | Hind | Middle | Front |
| $T_{sw}$ | Swing duration | 0.4 s - 1.6 s | | |
| $DS_{st}$ | Stance duty cycle | 0.6 | | |
| $h_{st}$ | Stance height | 0.07 m | 0.05 m | 0.09 m |
| $Tr_{body}$ | Body translation per step | 0.085 m | | |
| $h_{body}$ | Body height | Model:<br>0.175 m | Model:<br>0.175 m | Model:<br>0.175 m |
|  |  | Robot:<br>0.175 m | Robot:<br>0.19 m | Robot:<br>0.175 m |

#### 1) Inverse kinematics

Once a full foot trajectory has been generated, the solver performs inverse kinematics to find the joint angles necessary to complete the desired motions across the step. We chose to express the motion of the foot as a series of velocities in the body’s frame of reference, used each leg’s manipulator Jacobian to calculate the resulting joint angular velocities, then integrated these velocities to generate a trajectory of planned joint angle commands for the servomotors. This process was previously described in [72]. Because the change in configuration in this method is a linear approximation, error accumulates along the trajectory and the end will not exactly match the beginning. To join these ends without discontinuous motions (which produce large impulses in the inverse dynamics calculations), the first and final 30 points are replaced by a cubic interpolation between the 16th and *n*-15th points, where n is the total number of points. This process is repeated for each leg in the organism.

If the organism is configured to include parallel elastic elements, the solver additionally calculates equilibrium angles for each joint using the full RoM over the step. These values are tuned to produce leg positions resembling those of a newly dead fly, as such positions are the result of purely passive joint forces.

Once the joint angles, joint spatial positions, and leg segment CoM positions are calculated for each leg and for each timestep using the above method, the magnitudes of the angular velocity,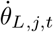 (where *L* is the leg number, *j* is the joint number, and *t* is the slice number out of *n* of the stepping cycle with duration Δ*t* = *T*_*sw*_*/n*) and angular acceleration,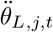 for each joint were calculated for each timestep as:

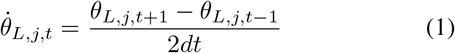

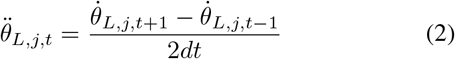

where dt is the amount of time in seconds each slice takes. If *t* = 1, the values for *t* = *n* are used in place of those for *t −* 1. Similarly, if *t* = *n* the values for *t* = 1 are used in place of those for *t* + 1.

Using these scalar values, the angular velocity vector for the CoM of each leg segment *s*, 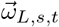is calculated as:

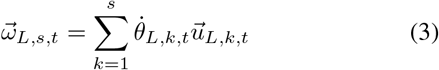

where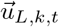 is the joint axis of joint *j* in leg *L* at slice *t*. For each leg segment, the corresponding *j* value is for the segment’s proximal joint. For example, *s* = 3 corresponds to the femur, while *j* = 3 corresponds to the TrF. This angular velocity value is then used to calculate the angular acceleration vector for each segment, 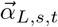:

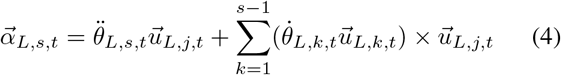

For the linear kinematics, the linear velocity of each segment’s CoM,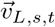, is calculated as:

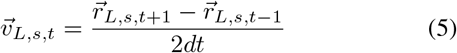

Where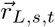 is the position of the segment’s CoM with respect to the origin of the spatial frame. The linear acceleration is similarly calculated as:

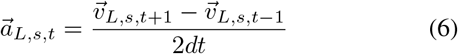

#### 2) Inverse dynamics

Finally, the solver performs inverse dynamic calculations to determine the forces 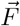and torques 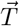acting on each leg segment and the thorax for each timestep. The solver shifts the phase of several of the joint angle arrays to produce the desired gait (e.g. for a tripod, 180 deg shift in one front leg, the contralateral middle leg, and the ipsilateral hind leg). If the organism includes parallel elastic components, the restoring torque for each timestep is calculated using the equilibrium positions defined during the inverse kinematic calculations.

The 3-D linear and angular Newtonian equations of motion (EoM) are then constructed for each segment of each leg, as well as the thorax. A Newtonian formulation was used in lieu of Lagrangian to be able to calculate internal reactions for each time step in addition to external forces. The equations are separated into x, y, and z components and assembled into a series of matrices in the form *Ax* = *b* for solving:

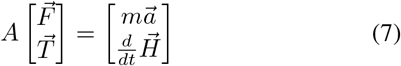

Where *m* is the mass of each segment, 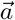 is the linear acceleration, 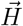is the angular momentum, and A is a matrix containing all of the coefficients of the force and torque variables in the left-hand sides of the EoM. Each segment’s portion of A, which we refer to as “building-blocks,” takes the form:

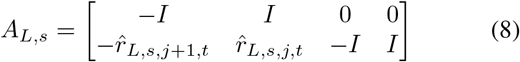

Where *I* is the 3×3 identity matrix and 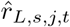 is the 3×3 skew matrix of the distance vector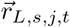 between the CoM of the segment and the segment’s proximal joint [70]. Similarly, 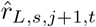is the 3×3 skew matrix of the distance vector between the CoM of the segment and the segment’s distal joint. The robot has 33 rigid bodies and 22 hinge joints, so assembling these building-blocks together results in a 192×210 matrix (i.e., 192 EoM and 210 unknown forces and torques). To ensure appropriate ground reaction forces during swing, we additionally add 3*L*_*st*_ equations to A, where *L*_*st*_ is the number of feet in swing, with the foot force of legs in swing explicitly set to zero. The thorax was assumed to only be moving at constant forward velocity to simplify calculations. With this number of equations and unknowns, the problem is underdetermined, and the forces and moments must be calculated by minimizing some objective function subject to the constraint that the EoM are satisfied. Our objective function represents the complementary potential energy *V* ^*∗*^ of the system [73]:

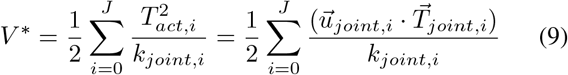

In this equation, *T*_*act*_ is the torque output for each actuated joint, *k*_*joint*_ is the stiffness in the joint from the actuator and any optional parallel elastic components, 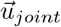 is the joint axis for the selected joint, and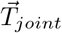 is the moment at the joint in the spatial frame. The stiffness in our chosen family of actuators, the Dynamixel MX series servos, was characterized for these calculations based on the process presented in [74]. By summing these values for the total number of actuated joints, *J*, created an equation that reflects both the actuator torques and the inherent elasticity in the system. Minimizing this value then minimizes the actuator torque while considering these other values. The program then minimizes Eq. 9 using the MATLAB function fmincon(), using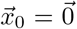 as the start point and the EoM as linear constraints. Once values are found, *T*_*act*_ is calculated by taking the dot product of each joint’s torque, 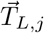, and 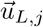. This process is repeated for every slice of the stepping period.

The final output of our solver is joint angles, velocities, and accelerations, leg segment and thorax internal forces, joint torques, and GRF for each timestep throughout a single step cycle. These data can then be concatenated to produce data for multiple steps. The joint angle data can be sent to the robot as commands for the servomotors. The dynamics data enabled us to carefully design the physical robot’s mechanical strength, i.e., the torque each actuator must deliver during walking.

### C. Comparison model generation

In order to validate the biomimicry of Drosophibot II, we created an additional *Drosophila*-like model to run through our solver framework. This model, called, “scale-*Drosophila*,” is a to-scale model of the fly based on the leg and thorax dimensions calculated as part of the kinematic analysis in Section II-A and presented in Table II. In the robot, some dimensions were altered from the animal kinematics for mechatronic feasibility (see section II-D for more details); these dimensions were retained at their measured values for scale-*Drosophila*. Additionally, each ThC joint was given the mobility observed in the animal, resulting in six total DoF per leg. The mass of this model was also distributed in a more animal-like way. In *Drosophila*, the abdomen, head, and thorax make up the bulk of the insect’s mass, with the legs only comprising around 11% of the total [34]. By contrast, the weight of Drosophibot II’s actuators results in 27% of the total weight distributed throughout the legs. To match the insect distribution in scale-*Drosophila*, we distributed the mass of a female fly (as reported in ref. [59]) using the percentages in ref. [34]. The CoM of the thorax, head, and abdomen were set at the same location as found in ref. [34]. The 11% of mass allocated to the legs was distributed throughout the leg segments proportional to each segment’s length. These considerations result in scale-*Drosophila* being a fully proportional fly analog. As force data from the animal is not presently available, this model allowed us to approximate force data and directly compare the dynamics of the animal and robot.

**TABLE II.** Leg segment lengths of Drosophibot II and *Drosophila. Drosophila* segment lengths were averaged from fly 3D motion capture data then scaled such that the middle leg femur was 100mm. Dimensions from Drosophibot II marked with * were lengthened due to mechatronic constraints.

| Segment | Drosophibot II Value (mm) |  |  | Scaled *Drosophila* Value (mm) |  |  | Percent Error |  |  |
| --- | --- | --- | --- | --- | --- | --- | --- | --- | --- |
|  | Hind | Middle | Front | Hind | Middle | Front | Hind | Middle | Front |
| Coxa | - | - | 54.50 | 19.96 | 12.77 | 49.34 | - | - | 10.46 |
| Trochanter | 27.97* | 27.97* | 27.97* | 9.684 | 9.851 | 6.372 | 188.8 | 183.9 | 339.0 |
| Femur | 100.4 | 100.4 | 80.00 | 95.83 | 100.0 | 78.43 | 4.769 | .4000 | 2.041 |
| Tibia | 97.85 | 97.85 | 77.84 | 94.43 | 90.48 | 67.45 | 3.622 | 8.145 | 15.40 |
| Tarsus | 120.0 | 100.0 | 69.60 | 119.0 | 99.96 | 79.33 | .8403 | .0400 | 12.27 |

### D. Robot construction

Drosophibot II is a 22 degree of freedom hexapod robot developed using our inverse kinematic and dynamic solver (Fig. 1A). Each leg is segmented similarly to the leg segments of *Drosophila*, with a varying number of joints actuated within each leg pair. The middle and hind pairs of legs have three DoF, corresponding to the CTr, TrF, and FTi joints (Fig. 1(B)). The front legs have an additional two DoF for the ThC1 and ThC3 to enable the more complex motions *Drosophila*’s front limbs undergo during walking (Fig. 1(C)). DoF that were omitted from the robot’s legs were fixed at their average position during the kinematic analysis [see previous sections and [65]]. Each joint is actuated by a Dynamixel MX-series smart servo (Robotis, Seoul, South Korea); MX-28s are used for most joints, whie MX-64s are used for the CTr due to the increased torque requirement at this joint. The majority of Drosphibot II’s components are 3-D printed out of Onyx composite nylon (Markforged, Waltham, MA). The distal tip of each tarsus is a 3-D printed Onyx core coated with a layer of Dragon Skin 10 silicone rubber (Smooth-On Inc., Macungie, PA). The silicone increases the robot’s traction on the substrate. These tips can be unthreaded from the tarsus and swapped for other designs to best suit the robot’s present terrain.

The robot’s leg segments were designed such that their relative proportions were similar to those of *Drosophila* with the length of the femur set to 10cm. This constraint results in the robot being approximately 140:1 scale to the insect. Table II compares leg segment lengths of Drosophibot II with the scaled-up average dimensions of the *Drosophila* specimens recorded as part of our kinematic analysis. In general, Drosophibot II’s leg segment lengths are within 16% of a 140:1 scale *Drosophila*. The lengths that are not within these margins had to be modified due to mechanical constraints. For example, the trochanter was lengthened to the minimum distance necessary for the bracket to rotate around the body of the servo. The lengths that were modified are indicated by asterisks (*) in Table II. Each leg was affixed to the thorax such that the CTr joints were approximately within the same horizontal plane.

Because 45% of *Drosophila*’s mass is in its abdomen, the CoM of Drosophibot II’s thorax and legs is much farther forward than in the insect. Previous investigations have calculated the anterior-posterior location of the insect’s CoM as between the middle and hind legs [34], while Drosophibot II’s CoM naturally falls between the middle and front legs. To shift the CoM to a more biological location, we have included an abdominal segment on Drosophibot II with slots for additional weight. We have found that including 1kg of mass approximately 125mm from the end of the thorax is enough to shift the CoM to an animal-like position. While adding mass does increase the load on the actuators, particularly in the hind legs, the robot’s strength-to-weight ratio is such that the required joint torques are still well within the operating limit of the servos. This weight is presently provided by a 1kg lab weight, but could be replaced by more functional ballast (e.g., batteries, sensors, control boards) in the future.

The robot is powered by an external Mean Well HEP-600-12 power supply (Mean Well Enterprises Co., Ltd., New Taipei City, Taiwan) able to supply up to 40 A at 12 V. Power is routed from the supply to each servo by a custom circuit board. This board also includes communication traces to the servos. Presently, control signals are provided by an external laptop through a U2D2 serial converter (Robotis, Seoul, South Korea) connected to the power board. The laptop runs a MATLAB script that writes servo angle commands formulated by the kinematic solver to each servo. The script then reads the present servo angles, as well as the current draw of each servo. This current draw is converted into torque based on ratios presented in the Dynamixel E-Manuals. The read-write loop runs at 65 Hz.

### E. Walking Experiments

To validate the performance of Drosophibot II after manufacture and assembly, we conducted experiments in which the robot walking forward and backward in a straight-line. Experiments were conducted on a sand-paper substrate in the Neuromechanical Intelligence Lab at West Virginia University in Morgantown, WV. The robot was sent joint angle commands from our inverse kinematic solver and returned the actual joint angles and the measured current (as an approximation of joint torque) for joint torque back to the control computer. It is presently unclear how accurate these current measurements are for the ground-truth, instant-to-instant joint torque. However, they are sufficient to provide general comparisons between different stepping scenarios of the physical robot.

Some parameters, such as floor level, were changed from those used solely in the solver to account for differences in rigidity (e.g., gear backlash, 3-D printed material elasticity, foot slippage) causing unwanted deflections and sagging. These changes brought the posture of the robot closer to the planned posture, and are presented in Table I.

During these walking experiments, the robot was commanded to step at periods between 1s - 4s. This range of walking speed was selected to dynamically scale the motions of the robot to those of the insect. Due to the robot’s much larger size, the robot will naturally have a different balance of viscous, elastic, and inertial forces present during stance and swing than *Drosophila*. Changing the walking speed can shift these values into a regime more similar to the animal. Detailed dynamic scaling calculations are presented in Appendix B.

## III. Results

Videos S1 and S2, provided in the supplementary material at http://ieeexplore.ieee.org, show the robot walking forward and backward, respectively. The following sections discuss the robot’s performance in simulation and in physical walking in greater detail.

### A. Robot validation in simulation

In order to evaluate the biomimicry of our robot, we kinematically and dynamically compared Drosophibot II to our scale-*Drosophila* model in our solver. For these calculations, the modeled robot walked with a period of 4s, with a stance duty cycle of 0.6 (full stepping parameters can be found in Table I). The chosen parameters produce interleg coordination used by flies at intermediate walking speeds, which is neither a pure tripod nor tetrapod [34], [59].

#### 1) Kinematic Comparison

Figure 4 compares the kinematics of each mobile DoF in a hind, middle, and front leg of Drosophibot II (solid lines) and scale-*Drosophila* (dashed lines) during a single step using the same footpaths scaled to each organism’s size. To aid in comparison, DoF that perform similar motions in the leg (e.g., both the ThC2 and CTr levate/depress the leg) are grouped on the same plot. Including all biological DoF in a leg predictably changes the motions of each individual DoF during walking between the organisms. In some cases, the change in angles in DoF present across both organisms is negligibly small. The FTi angles in the hind and middle legs are within 0.1 rad of each other throughout the step (Fig. 4G-I). The ThC1 and CTr angle in the front legs (Fig. 4C,F, respectively) show similarly small variation. In shared DoF with larger discrepancies, the majority exhibit similar trajectories and RoM over the step; the mid-point of that RoM is simply shifted. The ThC3 and TrF angles in the front limbs (Fig. 4C), CTr angles in the middle limbs (Fig. 4E), and FTi angles in the front limbs (Fig. 4I) follow this trend.

**Fig. 4.**
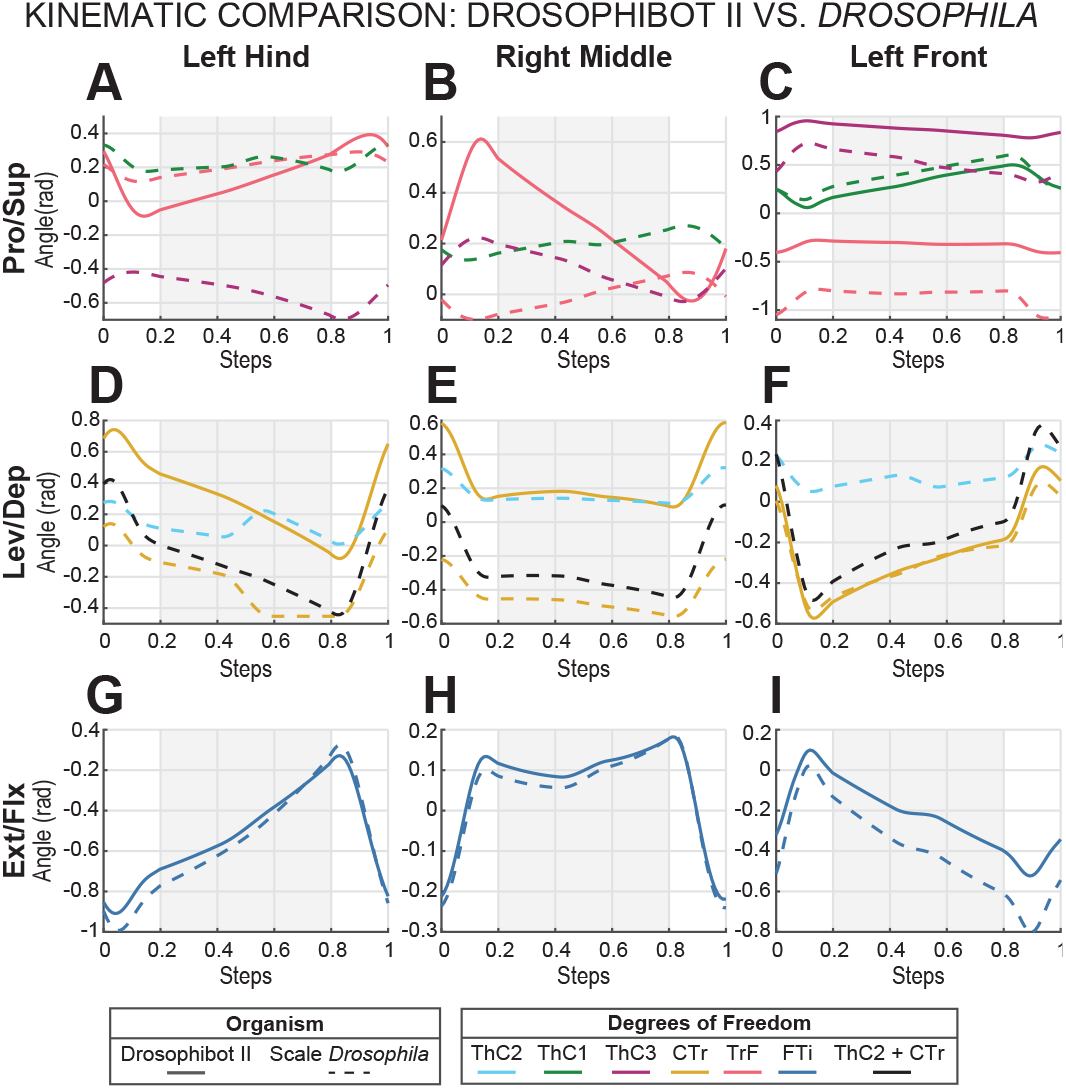
Modeled angles for each mobile DoF in the Drosophibot II (solid) and scale-*Drosophila* (dashed) organism models throughout a step for a hind (A,D,G), middle (B,E,H), and front (C,F,I) leg. DoF are grouped by the type of motion they provide in the leg: pronation/supination (A-C), levation/depression (D-F), and extension/flexion (G-I). Grey shaded areas represent the stance phase of the leg. The sum of the ThC2 and CTr angles from the scale-*Drosophila* model is additionally provided in black in (D-F).

We additionally found that in cases where multiple DoF acted in the same plane, such as the levation/depression DoF, summing the angles of scale-*Drosophila*’s multiple DoF produces a profile similar to that of the single DoF in Drosophibot II.To illustrate this point, the sum of the ThC2 and CTr angles of scale-*Drosophila* is presented in black in Figure 4D-F. For the middle and hind limbs (Fig. 4D-E), the combination of the ThC2 and CTr have trajectories and RoM more similar to Drosophibot II’s CTr angles than scale-*Drosophila*’s CTr angles alone.

The only DoF that does not align with these trends is the middle leg TrF (Fig. 4B). This DoF in Drosophibot II has a much larger range of motion than in scale-*Drosophila*, and the slope of the curves are opposite throughout the step. The profile of the robot motions more resembles that of the insect model’s ThC3. Both of these DoF provide pronation/supination through a rolling motion around the distal segment’s major axis in our models.

Overall, the angles of 20 of the 22 DoF in Drosophibot II either closely mapped to scale-*Drosophila*’s shared DoF, were of a similar trajectory and RoM to the shared DoF, or were of similar profile and RoM to the sum of all DoF providing that type of motion in the animal model. This demonstrates a capability of our robot to produce fly-like movements in individual leg segments.

#### 2) Dynamic Comparison

Figure 5 presents a comparison between Drosophibot II (solid lines) and scale-*Drosophila* (dashed lines) of the torques required from each DoF over a single step to produce the kinematics presented in Figure 4. Two different cases were considered; one in which no parallel stiffness was added to the joints (Fig. 5A), and one in which parallel elastic components were included (Fig. 5B). Elastic and viscous forces from the muscles and other tissues within the leg have been found to dominate motions in insects [61], [60], so analyzing these two cases allowed us to consider the effects of elasticity on both organisms’ dynamics. We used a joint stiffness value of *k*_*joint*_ = 3 *×* 10^*−*9^ Nm/rad for scale-*Drosophila*, and a value of *k*_*joint*_ = 1 Nm/rad for Drosophibot II. The value for *Drosophila* was calculated based on dynamic scaling calculations presented in ref. [62]. The value for Drosophibot II was selected such that the resulting torques at each DoF would not overload our selected actuators and the stiffness was feasible to produce with physical components at the scale of the robot. To aid in comparison, the torques calculated for scale-*Drosophila* were scaled by a factor of 5 *×* 10^8^.

**Fig. 5.**
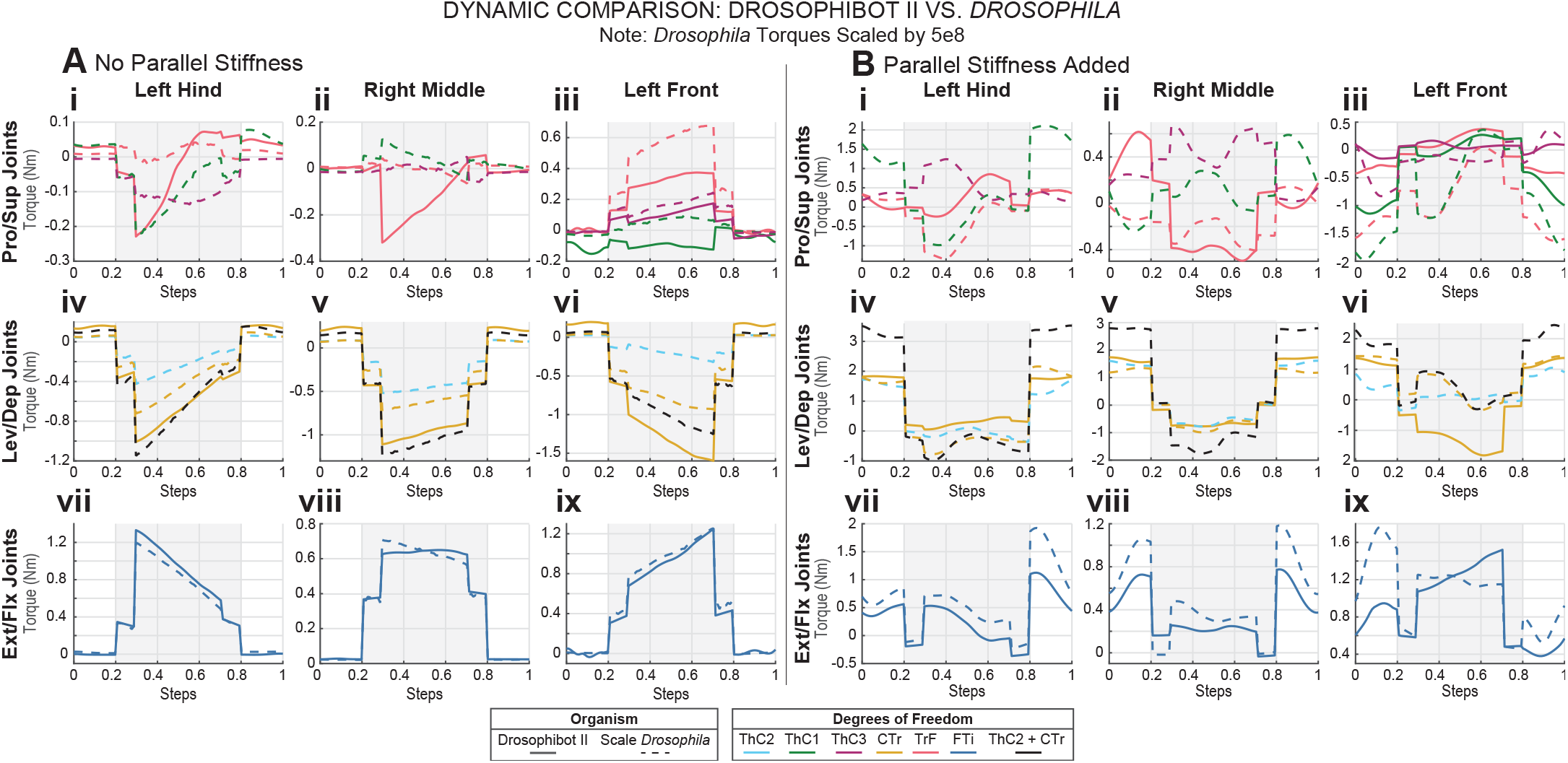
Modeled torques needed from each DoF in the hind (Ai,iv,vii; Bi,iv,vii), middle (Aii,v,viii; Bii,v,viii), and front (Aiii,vi,ix; Biii,vi,ix) Drosophibot II (solid) and scale-*Drosophila* (dashed) organism models to complete a step with the kinematics presented in Figure 4. Joints are grouped by the type of motion they broadly provide in the leg. Grey shaded areas represent the stance phase of the leg. The sum of the ThC2 and CTr angles from the scale-*Drosophila* model is additionally provided in black. scale-*Drosophila* torques were scaled by a factor of 5e8 to aid comparison. (A) Torque comparison of both models with no parallel stiffness added to the joints. (B) Torque comparison of both models with a parallel stiffness of 1 Nm/rad in Drosophibot II and 3e-9 Nm/rad in scale-*Drosophila*. Equilibrium positions for each joint were based on the resting posture of dead flies.

Without parallel stiffness, many of the DoF’s dynamics follow the trends outlined for the kinematics. Drosophibot II’s torques in the FTi joints across all legs (Fig. 5Avii-ix) and the ThC3 in the front legs (Fig. 5Aiii) closely agree with of those in scale-*Drosophila*. Additionally, the summation of the torques in the ThC2 and CTr of scale-*Drosophila* similarly agree with the robot’s CTr torques in the middle and hind limbs (Fig. 5Aiv-v). The front limb CTr torques (Fig. 5Avi), as well as the hind and front TrFs (Fig. 5Ai, Aiii) have more of a discrepancy between the two organisms. However, the trajectories of the torques over the step are still similar.

Also similar to the kinematics data, the middle TrF torque (Fig. 5Aii) exhibits unique behaviors from the rest of the DoF. Its magnitudes are much larger in the robot model than the insect model, and the profile is of opposite slope throughout the step. Unlike the kinematics, the torque curve does not instead resemble that of the ThC3 in scale-*Drosophila*.

Including elastic elements in parallel with the joints (Fig. 5B) produces substantial effects on the torques required from each DoF. The largest magnitude torques now occur during swing instead of stance, as the most dominant force in the leg becomes the restoring moments from the elastic components. The force of supporting the body helps resist these moments, lowering the torque during this phase of the step. Throughout the step, the torques in shared DoF no longer agree closely in magnitude in most cases. However, the trajectories of these pairs of curves remain similar. This similarity is particularly notable in the middle TrF (Fig. 5Bii), as this DoF had opposite trajectories in the kinematics and dynamics without parallel stiffness. During swing phase, the profiles are still of opposite slope. However, during stance the two curves follow a similar trajectory.

Unlike in the previous cases, the torque trajectories from Drosophibot II’s CTr more closely resemble those of scale-*Drosophila*’s CTr alone than the summation of the ThC2 and CTr (Fig. 5Biv-vi). In the middle and hind legs, both curves follow a similar profile, with a slight shift in overall magnitude. In the front legs, the trajectories of both curves closely match in swing. However, during the transitions between six legs in stance (0.2s-0.3s, 0.7s-0.8s) and three legs in stance (0.3s-0.7s) the torques move in opposite directions; Drosophibot II CTr torques decrease while the scale-*Drosophila* CTr torques increase. During the tripod portion of stance, the two curves follow similar trajectories.

Overall, 18 of the 22 DoF of Drosophibot II produce torques of similar trajectories over the entire step as our model of *Drosophila*, and 20 of 22 DoF produce a similar profile across stance phase.

### B. Effects of mass distribution

Non-biological distribution of mass is a common critique of biomimetic robots such a Drosophibot II, as increased mass in the legs could result in substantially different moments of inertia, affecting walking dynamics [60]. To address these concerns, we modeled the inverse dynamics of the robot in our solver, and ran tests with the physical robot, for multiple stepping periods and observed how the joint torques changed with stepping period. Figure 6 shows the modeled torques for each DoF in Drosophibot II, as well as the mean estimated torques from the robot’s actuators over five steps, for a hind (Fig. 6A-B), middle (Fig. 6C-D), and front (Fig. 6E-F) leg at stepping periods of 1s (solid), 2s (dashed), and 4s (dotted).

**Fig. 6.**
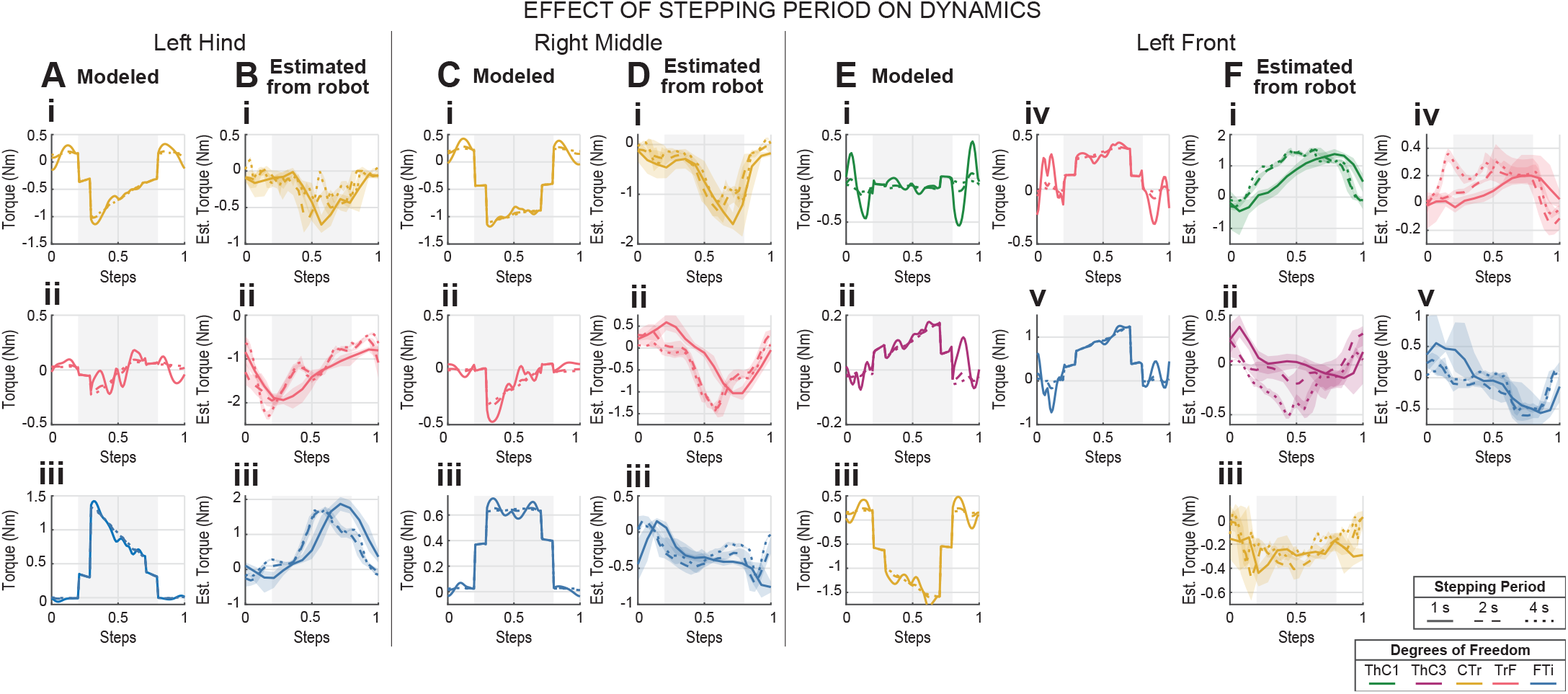
Joint torques for each joint in a hind (A-B), middle (C-D), and front (E-F) leg of Drosophibot II for stepping periods of 1s (solid), 2s (dashed), and 4s (dotted). Torques were both modeled in our solver (A,C,E) and estimated from current readings from the physical robot’s servos (B,D,F). Stepping data from the physical robot was averaged over five steps to generate a ‘typical’ step. Grey shaded areas represent the stance phase of the leg.

In simulation, decreasing the stepping period from 4s to 2s minimally affected the mean torques in each DoF (Fig. 6A,C,E). Changes in torque became most pronounced when decreasing to 1s steps, particularly in the swing phase of the front limbs, where torques increase by as much as 0.4 Nm (Fig. 6E). These dramatic increases are primarily during swing in the most proximal DoF, which must accelerate the full mass of the legs during this time.

As seen in the model, actuation torque in the proximal leg joints increases with stepping frequency in the physical robot data (Fig. 6B,D,F). While there is some slight deviation between the 4s and 2s stepping, the overall magnitude and phasing of the mean torques remains similar across all DoF. The greatest change occurs when the stepping period is decreased to 1s. This change most consistently results in a phase-lag of 0.1-0.175s compared to the 2s and 4s torques. There are additional effects on the mean torque magnitudes, but these effects are inconsistent between DoF. Some DoF experience an increase in the mean torque (Fig. 6Di), others experience a decrease (Fig. 6Bii), and still others are roughly unchanged (Fig. 6Fi) However, in most DoF this change in magnitude, if any, is within the range of variance between the steps used to form the mean value (shaded regions). The minimal changes in torque between the 4s and 2s cases in both the model and the physical robot demonstrates it is quasi-static in that range of speeds. As such, we slowed the robot’s behavior to minimize the effects of the non-biological distribution of mass on the dynamics of the robot in those range of walking speeds. Decreasing the stepping period to 1s begins to shift the robot into a more inertially-dominated regime, providing a functional upper limit of walking speed for our experiments. These experiments also highlight the ability of the solver to predict aspects of the physical robot.

### C. Effect of walking direction

We also used the physical robot to test how walking direction affects the dynamics in each DoF. Figure 7 compares the torques for each DoF averaged over five steps for each leg during forward (dashed) and backward (solid) walking at a 2s stepping period. To generate backward walking, the kinematics from forward walking were run in reverse. As such, *Tr*_*body*_ = *−* 0.085m. No other stepping parameters were modified, and are given in Table I.

**Fig. 7.**
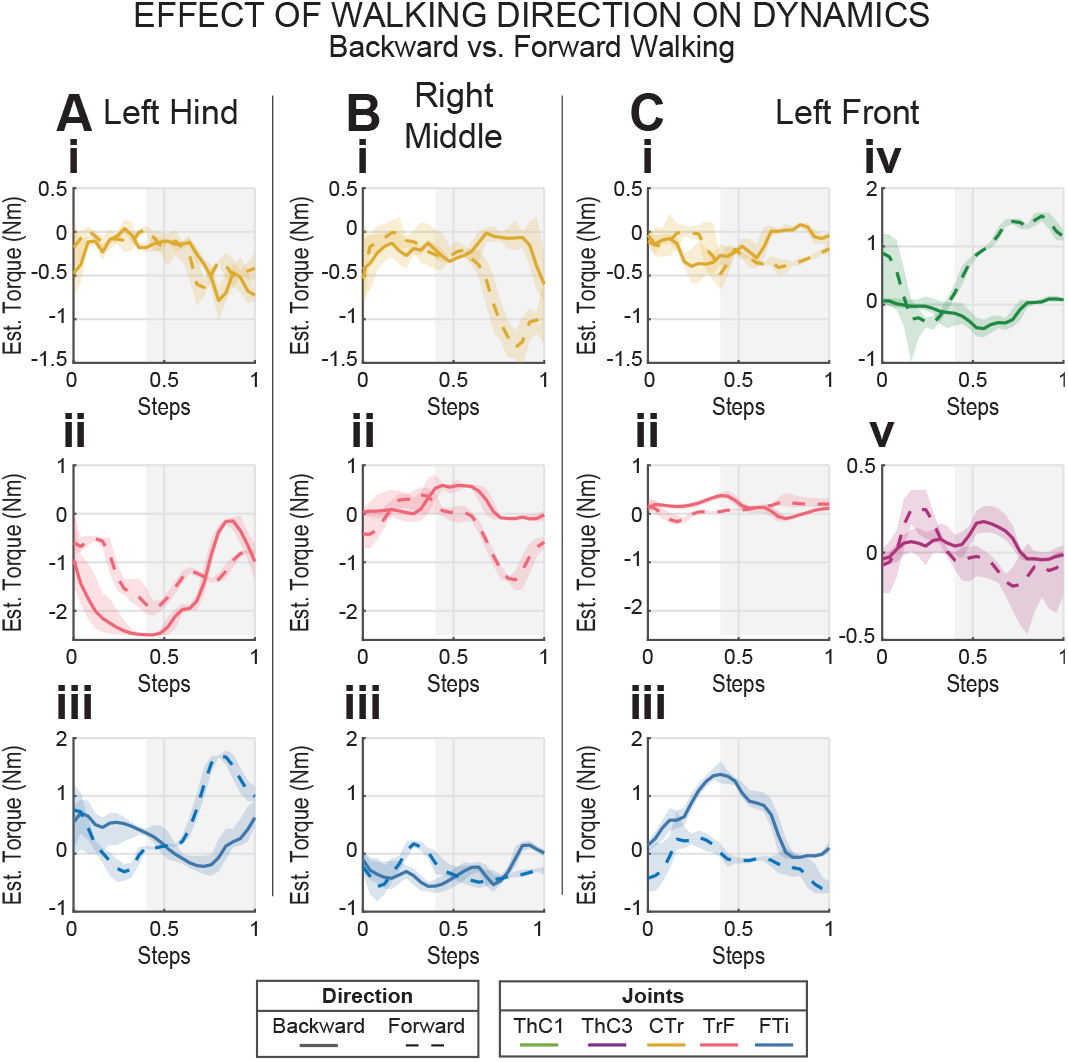
Comparison of joint torques estimated using current draw from each joint’s actuator in a hind (A), middle (B), and front (C) leg of Drosophibot II during forward (dashed) and backward (solid) walking at a 2s stepping period. To generate backward walking, kinematics from forward walking were run in reverse. Grey shaded areas represent the stance phase of the leg.

Reversing the walking direction changes the polarity and magnitude of several DoF’s torques in each leg pair. Which DoF are most affected varies across leg pairs. In the hind legs, walking backward increases the torque requirements of the TrF (Fig. 7Aii) and reverses the torque directions throughout the trajectory of the FTi (Fig. 7Aiii). In the middle legs, the magnitudes of the CTr (Fig. 7Bi) and the TrF (Fig. 7Bii) torques are decreased during stance, and the slopes are reversed such that the values approach zero or become positive, respectively, rather than remaining negative. The front leg ThC1 (Fig. 7Civ) and FTi (Fig. 7Ciii) undergo significant changes in magnitude during stance. Additionally, the trajectories of the front leg CTr (Fig. 7Ci) and ThC3 (Fig. 7Cv) change from concave-up to concave-down. Of the 22 DoF in Drosophibot, 14 experience significant changes to their magnitude or polarity when walking backward. However, these changed values are still within the capabilities of the actuators, and the robot is able to walk in either direction without issue.

## IV. Discussion

The development and experimental validation of Drosophibot II, a biomimetic robot modeled after an adult fruit fly, is presented. The robot was designed with close consideration of leg proportions and necessary DoF informed by walking kinematic data from the animals. Its mechatronics were designed using a program that calculates the inverse kinematics for biomimetic walking and solves the inverse dynamics throughout a single step. Using this solver program, Drosophibot II was compared to a scale model of *Drosophila*. The solver demonstrates that the two organisms produce similar motions in their leg segments while stepping through the same footpaths; the majority of robot DoF either had similar trajectories and ranges of motion to shared insect DoF, or to the sum of all insect DoF acting within the same plane. Dynamically, the majority of modeled torques for each robot DoF had similar trajectories to those of the insect in cases with and without insect-like parallel elasticity in the joints. Robot torques were also found to be similar across three different stepping periods, demonstrating that Drosophibot II is quasistatic for our intended walking speeds. Finally, the robot was able to execute straight-line forward and backward walking using the joint angles generated by our solver.

Completing mechatronic development of Drosophibot II enables a host of potential applications to neuromechanical research. The robot can be used as a test platform for nervous system-inspired walking controllers, also called synthetic nervous systems (SNS) [20], [53], [54], [55], such as the one previously developed for the first Drosophibot [24]. By modeling the dynamics of neurons and synapses, as well as the morphology of the nervous system, these controllers can serve to increase our understanding of how the nervous system controls legged locomotion and motor control in general. For example, many recent studies focused on how descending control contributes to the initiation, maintenance, and task-specific modulation of locomotive behaviors in *Drosophila* leading to identification of brain neurons involved in forward walking, turning, and backwards walking [47], [48], [49]. In this context, Drosophibot II can serve to test diverse putative descending control regimes based on data from neurobiological studies.

In regards to neuromechanical modeling, our inverse dynamic torque data for the robot and fly body-plans highlights the importance of considering joint elasticity in biomimetic platforms. Adding in insect-like parallel elasticity to our model significantly changed the torque magnitudes and trajectories for each part of a step in every DoF by producing elastic restoring moments. In other studies, such elastic components have been found to require different control mechanisms than those for inertially-dominated organisms, such as consistent motor-neuron activation during swing phase [60], [61], and “braking” forces from opposing muscles to tune/halt rapid passive restoring moments [75]. *Drosophila*’s nervous system has evolved to control body mechanics more resembling our model with parallel elasticity than the model without, so including parallel elastic components on Drosophibot II will be important for future endeavors into matching the mechanical and neural dynamics of biomimetic walking.

The robot can also serve as a data-collection platform for biomimetic sensory feedback. Several studies have previously used biologically-inspired robots to investigate sensory discharge from campaniform sensilla (CS), mechanoreceptors on the insect exoskeleton that encode strain [24], [76], [77], [78]. These studies primarily focus on single-leg stepping. By including strain gauges in biological locations on Drosophibot II’s limbs, strain data that may be available to the nervous system throughout six legged stepping can be hypothesized. To this end, it is beneficial that the torque trajectories modeled for the CTr and FTi in each leg were minimally affected by changing the body plan. The roles of CS fields on the trochanter and tibia have been investigated for many years in a variety of insects [79], [80], [81], [82], but recording from many sensilla in a walking animal is prohibitively difficult. Recording strain data from these locations on Drosophibot II will contribute to this body of work by producing strain readings from all CS locations on all legs at once that are functionally comparable to the animal.

Our solver program can also be used to analyze the mechanics of other animals and robots. While we have primarily used it thus far to validate the robot, any body plan or branching chain of rigid bodies can be input into the solver. As such, it could be used to design biomimetic robots based on any particular species, provided that motion capture data were available. Furthermore, because forces and torques are calculated using Newton-Euler equations of motion instead of Lagrangian, the reaction forces and moments at each joint are calculated. This means, for example, our *Drosophila*-scale body plan can be used generate theoretical values for animal-like GRFs and reaction forces and moments at each joint. The latter could be of particular interest for calculating biologically plausible limb stresses and strains for finite element analysis of the insect cuticle [82], improving understanding of what stresses and strains are experienced during locomotion, and how they activate CS across the leg.

Though we have carefully considered the kinematics and dynamics of Drosophibot II in order to match them to *Drosophila*, the present study does have limitations. First, although the leg segment lengths of the robot are closely proportional to the animal’s, the distances between the ThC joints (or where the ThC would be, in the case of the middle and hind legs) have significantly more error due to the size of the actuators. The actuators in Drosophibot II were positioned as close as possible to each other both laterally and along the body such that the distances between leg pairs were still proportional. For example, the lateral distance between the middle leg pair ThC joints was lengthened to the minimum distance possible without the CTr servos colliding, then the lateral distances between the front and hind limbs were also extended to keep the distances between the ThC pairs scaled biomimetically. However, the relative size of the actuators still produced thorax dimensions up to x10 greater than those in the fly. Our solver produces footpaths based on the location of each leg’s ThC in the global frame, so we do not believe these increased thorax dimensions significantly affect the kinematics and dynamics between our modeled organisms. In the future, the legs could be positioned closer together by moving the actuators into the body and using belts or cables to drive the joints [83], [84], [85].

Additionally, our inverse dynamic solver makes several simplifications that could affect the model’s accuracy to the physical robot’s dynamics. Our solver does not include difficult-to-simulate parameters such as slip and elastic deformation of the 3D printed materials, which can cause sagging of the robot posture. The solver also does not account for accelerations of the main body segment, which naturally occurs as the animal’s thorax bobs or rotates during walking. On the physical robot, the thorax CoM will naturally shift slightly as different leg combinations enter and leave stance. It is presently difficult to quantitatively determine how these simplifications affect our model’s accuracy, as the robot actuators’ “torque” readings are extrapolated from the current draw of the motor. While the manufacturer provides conversions for these readings, they have been found to have idiosyncratic inaccuracy in additional bench-top experiments. As such, the readings are primarily useful only for comparing robot data to other robot data. Future work will involve both attempting to minimize the effect of these un-modeled forces (e.g., increasing the coefficient of friction between the tarsi tips and the ground) and exploring alternate ways to read accurate actuator torque data from the robot.

## Appendix A Biological data collection

We used Bolt-GAL4*>*UAS-CsChrimson *Drosophila melanogaster* flies [48] for all experiments. This allowed optogenetic initiation of sustained, fast forward walking. Flies were reared on a standard yeast-based medium [86] at 25°C and 65% humidity in a 12h:12h day:night cycle and experiments were performed with 3-8 days post-eclosion males and females. Prior to experiments, the animals were kept in fresh vials in which the food was soaked with 50 μL of a 100 mmol L-1 all-trans-Retinal solution (R2500; Sigma-Aldrich, RRID:SCR 008988) in the dark for at least three days.

To capture leg movements of walking fruit flies, the animals were tethered and placed on an air-supported spherical treadmill setup as previously described in detail in [56]. For optogenetic activation of CsChrimson, we used a red laser (658 nm) targeting the animal’s head. Walking behavior was recorded with six synchronized high-speed cameras (acA1300-200um, Basler AG, Ahrensburg, Germany) equipped with 50 mm lenses (LM50JC1MS, Kowa Optical Products Co. Ltd., Nagoya, Japan). Cameras were arranged such that either body side was recorded simultaneously by three cameras providing a front, side, and hind aspect (Fig. 2). A custom-built in infra-red LED ring (880 nm) was mounted above the setup to illuminate the scene. Videos were acquired at 400 Hz, a resolution of 896 by 540 pixels, and an exposure time of 500 μs.

Automated tracking of body parts in the videos was performed with the DeepLabCut (DLC) toolbox [57]. We tracked six features on each leg: the thorax-coxa (ThC) joint, coxa-trochanter (CTr) joint, trochanter-femur (TrF) joint, femur-tibia (FTi) joint, tibia-tarsus (TiTar) joint, and the tip of the tarsus. Additionally, we tracked features on the fly’s body and head: the posterior scutellum apex on the thorax, the wing hinges, and the antennae. For tracking, we trained three independent ResNet-50 networks for the front, side, and hind camera groups (i.e., a single network was trained for the same two camera views of the left and right body side). Training sets were generated by manually annotating short walking sequences of three male and three female flies (630 images for each network) and extended by using the default in-built augmentation algorithm of DLC. Tracked feature positions were visually validated and manually corrected if necessary.

For 3D reconstruction of feature positions, we calibrated the cameras with a custom-made checkerboard pattern (7 × 6 squares with size 399 μm x 399 μm per square) developed on a photographic slide (Gerstenberg Atelier fuür Visuelle Medien, Templin, Germany). At first, the intrinsic parameters and lens distortion coefficients (three radial and two tangential coefficients) for each camera were determined, followed by pairwise stereo calibrations of adjacent cameras to obtain the extrinsic camera parameters. Average reprojection errors of less than one pixel were considered acceptable. For triangulation of 3D positions, a singular value decomposition algorithm was applied [87], [88]. 3D positions of the tracked features were transformed to a body-centered coordinate system derived from the triangle formed by the posterior scutellum apex and the right and left wing hinge.

All used custom-made devices were built by the Electronics workshop of the Institute of Zoology, University of Cologne. Custom-written functions for camera acquisition (Harvester library), camera calibration (OpenCV library), interacting with DLC, and 3D reconstruction were implemented in Python (version 3.95, RRID: SCR 008394).

## Appendix B Dynamic scaling

Two motions are dynamically scaled if they exhibit the same relative magnitude of different types of forces, e.g., inertial and elastic. The dimensionless parameter that describes the ratio between elastic and inertial forces with an oscillating system (e.g., a leg retracting and protracting) is the Strouhal number:

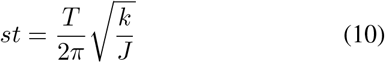

where *st* is the Strouhal number, T is the period of oscillation, k is the effective stiffness of a joint in units of Nm/rad, and J is the mass moment of inertia of the leg about the joint.

In previous work, the Strouhal number for the leg of a walking fruit fly was approximated to be 6.5 [24], indicating that the motion of the leg is dominated by elastic forces, not inertial forces. We experimentally determined the effective moment of inertia of the MX-64 servo motor that actuates the coxa-trochanter joint to be 0.016 *kgm*^2^. This number was several orders of magnitude larger than the moment of inertia due to the extended mass of the leg or the mass of the distal actuators. We experimentally determined the stiffness of the servo to be 20 Nm/rad due to the proportional feedback loop that controls its position. With an unknown stepping period *T*_*robot*_, the Strouhal number becomes:

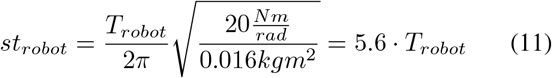

For the robot to have the same Strouhal number as the fly, *T*_*robot*_ = 1.16*s*. To add a safety factor and ensure the robot’s dynamics are not dominated by inertial forces, we run the robot anywhere between 1*s ≤ T*_*robot*_ *≤* 5*s*.

## Acknowledgment

The authors would like to thank Till Bockemuühl, Michael Duübbert, and Mehrdad Ghanbari for their help in developing the motion capture setup, as well as William Zyhowski for his help in developing the robot’s custom circuitry.

